# Optogenetic control of *Neisseria meningitidis* Cas9 genome editing using an engineered, light-switchable anti-CRISPR protein

**DOI:** 10.1101/858589

**Authors:** Mareike D. Hoffmann, Jan Mathony, Julius Upmeier zu Belzen, Zander Harteveld, Christina Stengl, Bruno E. Correia, Roland Eils, Dominik Niopek

## Abstract

Optogenetic control of CRISPR-Cas9 systems has significantly improved our ability to perform genome perturbations in living cells with high precision in time and space. As new Cas orthologues with advantageous properties are rapidly being discovered and engineered, the need for straightforward strategies to control their activity via exogenous stimuli persists. The Cas9 from *Neisseria meningitidis* (*Nme*) is a particularly small and target-specific Cas9 orthologue, and thus of high interest for *in vivo* genome editing applications.

Here, we report the first optogenetic tool to control *Nme*Cas9 activity in mammalian cells via an engineered, light-dependent anti-CRISPR (Acr) protein. Building on our previous Acr engineering work, we created hybrids between the *Nme*Cas9 inhibitor AcrIIC3 and the LOV2 blue light sensory domain from *Avena sativa*. Two AcrIIC3-LOV2 hybrids from our collection potently blocked *Nme*Cas9 activity in the dark, while permitting robust genome editing at various endogenous loci upon blue light irradiation. Structural analysis revealed that, within these hybrids, the LOV2 domain is located in striking proximity to the Cas9 binding surface. Together, our work demonstrates optogenetic regulation of a type II-C CRISPR effector and might suggest a new route for the design of optogenetic Acrs.

## INTRODUCTION

CRISPR (clustered regularly-interspaced short palindromic repeats)-Cas technologies facilitate site-specific targeting and manipulation of genes in living cells (1–3) and currently transform many areas of biomedical research. Class II CRISPR-Cas effectors are the major workhorses driving this transformation. They comprise only two parts, a Cas nuclease and a single guide RNA (sg)RNA, which directs the Cas nuclease to selected nucleic acid targets by means of sequence complementarity. Due to their simplicity and versatility, class II CRISPR systems enable a plethora of applications including targeted induction of DNA double-strand breaks for genome editing (2,4), regulation of endogenous transcription (4,5), epigenetic reprogramming (6–8), DNA labeling (9,10) and base editing (11,12).

The type II-A CRISPR-Cas9 from *Streptococcus pyogenes* (*Spy*Cas9) is the most widely applied CRISPR-Cas9 orthologue. Due to its large size of 1,368 amino acids (158 kDa) and high off-target rates (13–15), however, alternative CRISPR-Cas9 orthologues gained attention. A particularly interesting candidate is the type II-C Cas9 from *Neisseria meningitidis* (*Nme*Cas9). With only 1,081 amino acids (124 kDa), *Nme*Cas9 is considerably smaller than *Spy*Cas9. On top, *Nme*Cas9 exhibits an exceptionally high target specificity, possibly due its longer target recognition sequence (16,17). These properties render *Nme*Cas9 a powerful tool for various applications, including *in vivo* gene editing (18) and also RNA-induced genome binding via catalytically impaired *Nme*Cas9 mutants (19,20).

The ability to control and fine-tune *Nme*Cas9 activity via exogenous stimuli would further enhance the precision at which CRISPR genome perturbations can be made. Unlike *Spy*Cas9, for which a whole battery of tools exist that facilitate its conditional activation by chemical triggers (21–24), light (25–29) or temperature (30,31), no method for conditional activation of *Nme*Cas9 by exogenous triggers has yet been developed.

Anti-CRISPR (Acr) proteins are bacteriophage-derived antagonists of CRISPR-Cas systems (20,32–43). They represent a highly diverse class of proteins practically without structural and sequence homology to other proteins (38,39). Acrs inhibit Cas nucleases by various mechanisms, including inhibition of DNA binding (44,45), cleavage of sgRNAs (46), masking catalytic residues and/or inducing Cas9 dimerization (47). AcrIIC3 is an incredibly potent *Nme*Cas9 inhibitor initially discovered in a putative prophage element within the *Neisseria meningitidis* genome. It binds the catalytic HNH domain and induces dimerization of *Nme*Cas9 via the REC lobe in a ratio of AcrIIC3:Cas9 = 2:2, thereby blocking DNA binding (47,48).

Here, we report the engineering and application of CASANOVA-C3 (for **C**RISPR-Cas9 **a**ctivity **s**witching via **a n**ovel **o**ptogenetic **v**ariant of **A**crII**C3**), a light-dependent anti-CRISPR protein for conditional activation of *Nme*Cas9. Building on our recent AcrIIA4 engineering work (29,49), we created hybrids between AcrIIC3 and the *Avena sativa* (*As*)LOV2 photosensory domain by systematically sampling AcrIIC3 surface sites. Following screening and optimization, two AcrIIC3-LOV2 hybrid variants were obtained, which potently inhibit *Nme*Cas9 in the dark, while releasing its activity upon blue light irradiation (Figure 1A). We demonstrate light-dependent editing of various genomic loci upon transient transfection and Adeno-associated virus (AAV)-mediated transduction. Finally, using structural modeling, we show that our CASANOVA-C3 design presents an unconventional, yet potentially powerful blueprint to engineer light-dependent protein-protein interactions.

**Figure 1.**
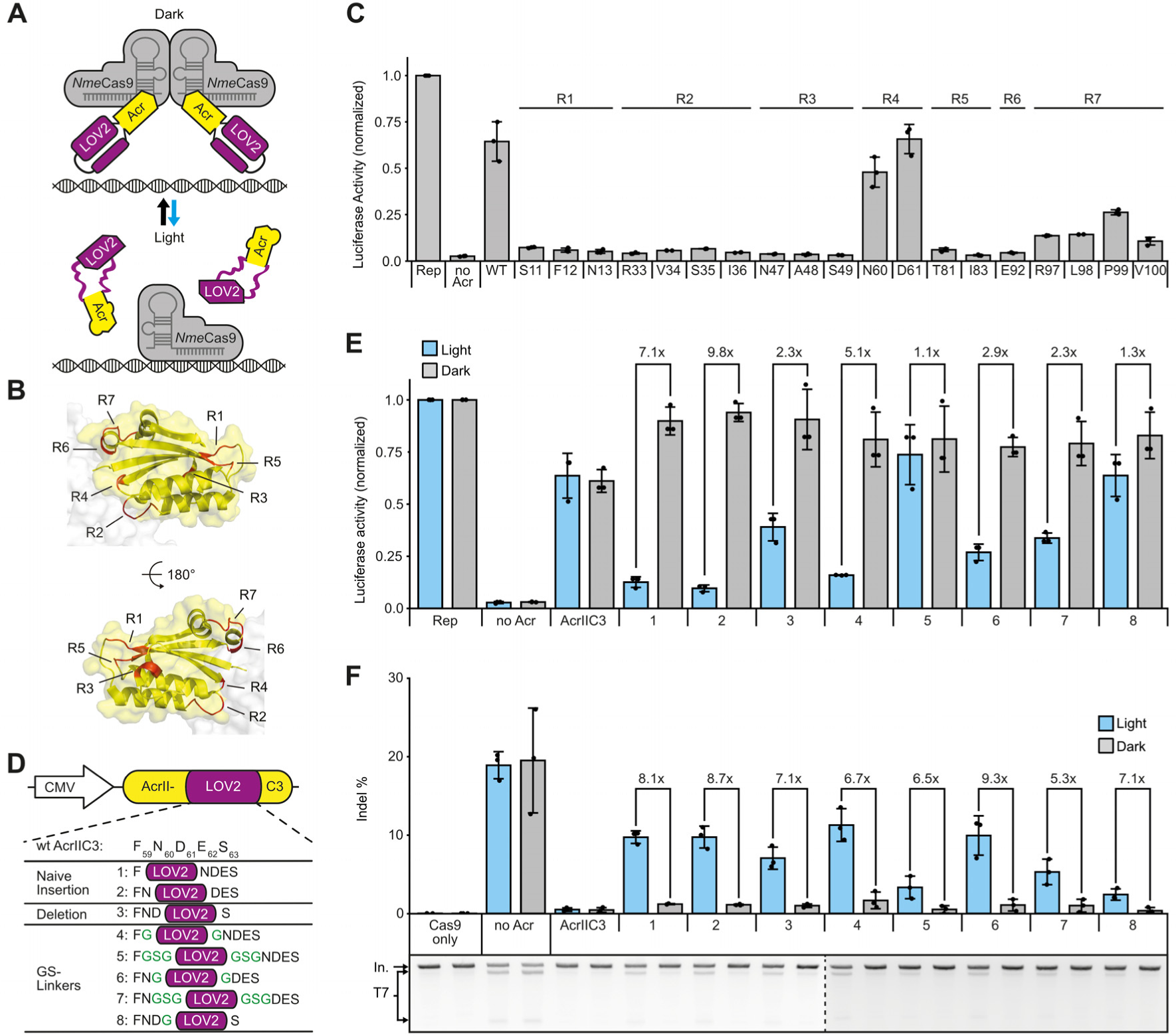
Engineering of CASANOVA-C3, a light-switchable anti-CRISPR protein for optogenetic control of *Nme*Cas9 (**A**) Schematic of CASANOVA-C3 function. (**B**) Structure of AcrIIC3. The seven regions chosen for LOV2 domain insertion (R1-R7) are shown in red (PDB 6J9N). (**C**) Luciferase reporter-based screen of AcrIIC3-LOV2 hybrids. HEK293T cells were co-transfected with vectors encoding (i) a firefly luciferase reporter, (ii) *Nme*Cas9 and a sgRNA targeting the luciferase reporter and (iii) either wt AcrIIC3 or the indicated AcrIIC3-LOV2 hybrid followed by luciferase assay. The AcrIIC3 residues behind which the LOV2 domain was inserted are indicated. R1-7 correspond to the different regions in **B.** R, region. Rep, reporter only control. (**D**) Lead panel of AcrIIC3-LOV2 hybrids. Glycine-serine linkers are in green. (**E**) Luciferase assay screen of the AcrIIC3-LOV2 hybrids in **D**. Cells were transfected as in **C** and then exposed to blue light or kept in the dark for 48 hours, followed by luciferase assay. Rep, reporter only control. (**F**) HEK293T cells were co-transfected with vectors encoding (i) *Nme*Cas9 and a sgRNA targeting the endogenous IL2RG locus and (ii) the indicated Acr variant in **D**. Samples were exposed to blue light or kept in the dark for 72 hours. Gene editing was assessed by T7 assay. Representative gel images are shown below the bar charts. The dotted line separates different gels. In, input. T7, T7 cleavage fragments. (**C, E, F**) Bars represent mean values, error bars the standard deviation and dots individual data points from n = 3 independent experiments.

## MATERIAL AND METHODS

### General methods and cloning

A list of all constructs created and used in this study is provided in Supplementary Table 1. Annotated plasmid sequences (SnapGene DNA files) are provided as Supplementary Data. Oligonucleotides and double-stranded DNA fragments were obtained from Integrated DNA Technologies. AAV plasmids were generated via restriction enzyme cloning, all other constructs were cloned by Golden Gate assembly (50). The vector for co-expression of *Nme*Cas9 and the VEGFA sgRNA was previously published by us (51). All other sgRNAs were cloned into plasmid pEJS654 All-in-One AAV-sgRNA-hNmeCas9 (kind gift from Erik Sontheimer, Addgene plasmid #112139) via the SapI restriction sites. A list of genomic target sites is provided in Supplementary Table 2. The dual luciferase reporter was previously reported by us (49). AcrIIC3-LOV2 hybrid constructs were created by inserting the LOV2 domain into our published CMV-driven AcrIIC3 expression vector (Addgene plasmid #120301) (51). To this end, the AcrIIC3 vector was linearized by an around-the-horn PCR using primers carrying BbsI restriction sites as 5’ extension. The LOV2 domain was PCR amplified from the vector CMV-CASANOVA (Addgene plasmid #113035) previously reported by us (29). LOV2 primer 5’ extensions contained BbsI sites compatible with the vector amplicon BbsI sites and – optionally – sequences encoding flexible, glycine-serine linkers. Golden-gate cloning was then used to assemble AcrIIC3-LOV2 hybrids. Point mutations for the LOV2-RVH variant were introduced by site-directed mutagenesis. AcrIIC3-LOV2 AAV vectors were created by replacing the wild-type AcrIIC3 coding sequence in vector AAV CMV-driven AcrIIC3-scaffold (2xBsmBI sites) previously reported by us (51) by the AcrIIC3-LOV2 coding sequences via the XhoI and NheI restriction sites.

All PCRs were performed using the Q5 Hot Start High-Fidelity Polymerase (New England Biolabs, NEB). PCR products were analysed on 1-2 % TAE or TBE agarose gels. The desired bands were cut out and extracted from the gel using a QIAquick Gel Extraction Kit (Qiagen). Restriction digests and Golden Gate assembly were performed according to the manufacturer’s protocols with enzymes obtained from Thermo Fisher Scientific or NEB. Fragments were ligated using T4 DNA Ligase (Thermo Fisher Scientific) and constructs transformed into chemically-competent Top10 cells (Thermo Fisher Scientific). Plasmid DNA was purified using QIAprep Spin Miniprep, Plasmid Plus Midi or Plasmid Maxi Kit (all Qiagen).

### Cell culture and AAV production

HEK293T (human embryonic kidney) cells were maintained in DMEM (Thermo Fisher Scientific) supplemented with 10 % (v/v) FCS (Biochrom AG), 2 mM L-glutamine (Thermo Fisher Scientific) and 100 U/ml penicillin and 100 μg/ml streptomycin (Thermo Fisher Scientific). Cells were cultured at 37 °C and 5 % CO_2_ in a humidified incubator and passaged every 2-3 days (i.e. when about 90 % confluent). Cells were authenticated and tested for mycoplasma contamination prior to use via a commercial service (Multiplexion).

To produce AAV lysates, low passage HEK293T cells were seeded at a density of 350,000 cells per well into six-well plates (CytoOne) using 4 ml of medium per well. The next day, cells were triple-transfected with (i) AAV vector (transgene flanked by AAV ITRs), (ii) the AAV helper plasmid carrying the *rep* and *cap* genes of AAV serotype 2 and (iii) an adenoviral plasmid providing helper functions for AAV production using 8 μl TurboFect Transfection Reagent per well (Thermo Fisher Scientific). 72 hours post transfection, cells were flushed off the culture plate surface by pipetting and collected into the medium. Samples were then spun down, the supernatant (medium) was discarded and the cell pellet was re-suspended in 300 μl PBS. Cells were then lysed by applying five freeze-thaw cycles of snap-freezing in liquid nitrogen, followed by incubation at 37 °C in a water bath. Subsequently, the cell lysate was centrifuged at 18,000 g at 4 °C for 10 min to remove cell debris and the AAV-containing supernatant was kept at 4 °C until use. Lysates were stored for no longer than 3 weeks.

### Blue light setup

Blue light exposure of the samples was achieved with a custom-made LED setup, consisting of six blue light high power LEDs (type CREE XP-E D5-15; emission peak ~460 nm; emission angle ~130°; LED-TECH.DE) connected to a Switching Mode Power Supply (Manson; HCS-3102). The setup was controlled by a custom Python script, running on a Raspberry Pi. LEDs were positioned underneath a transparent table made of acrylic glass and positioned inside a cell culture incubator. Culture plates with samples were positioned on top of the table, i.e. they were irradiated from below through the acrylic glass and culture plate’s transparent bottom. Illumination intensity was set to 3 W/m^2^ and regularly confirmed by measurements with a LI-COR LI-250A light meter. Pulsatile illumination was used (5 s on, 10 s off). Dark control samples were kept in the same incubator, but protected from light by covering the transparent sample plate parts with black vinyl foil (Starlab).

### Luciferase assay

12,500 cells/well were seeded into black, clear-bottom 96-well plates (Corning). The next day, cells were transfected using Lipofectamine 3000 Reagent (Thermo Fisher Scientific), following the manufacturer’s protocol. 200 ng of total DNA per well comprising equal amounts (plasmid mass) of (i) dual luciferase reporter plasmid, (ii) *Nme*Cas9 and sgRNA encoding plasmid and (iii) Acr-LOV2 constructs were co-transfected. The reporter construct encoded a *Renilla* and firefly luciferase as well as a sgRNA targeting a sequence stretch implanted in frame with (5’ of) the firefly coding sequence. 48 hours post transfection, cells were washed with 1x PBS (Sigma-Aldrich) and lysed in Passive Lysis Buffer (Promega). Subsequently, the luciferase activities were measured with the Dual-Glo Luciferase Assay System (Promega) on a GLOMAX discover or GLOMAX 96-microplate luminometer (both Promega). Integration times of 10 s were used; delay between automated substrate injection and measurement was 2 s. Firefly luciferase photon counts were normalized to *Renilla* luciferase photon counts. Finally, obtained values were normalized to the reporter only controls in the light or dark.

### T7 endonuclease I assay (T7 assay)

HEK293T cells were seeded into black, clear-bottom 96-well plates using 12,500 cells and 100 μl medium per well (Corning). The next day, cells were transfected with 150 ng total DNA per well using the Lipofectamine 3000 reagent and following the manufacturer’s protocol. The vector mass ratios of Cas9/sgRNA and AcrIIC3-LOV2 construct used during transfection are indicated in the corresponding figures.

For AAV experiments, 3,500 HEK293T cells were seeded per well into black, clear-bottom 96-well plates (Corning). The cells were co-transduced twice, i.e. on two consecutive days, with 80 μl AAV lysate. The lysate comprised Cas9/sgRNA and AcrIIC3-LOV2 AAV lysate in a volumetric ratio as indicated in the corresponding figures. For the controls without Acr, Cas9/sgRNA AAV lysate was topped up to 80 μl with PBS to keep the transduction volume constant between all samples. Cells were lysed 72 hours post transfection or post (first) transduction using the DirectPCR lysis reagent (PeqLab) supplemented with 200 μg/ml proteinase K (Roche Diagnostics). The targeted genomic locus was then PCR amplified with primers flanking the expected cutting site (Supplementary Table 3) using Q5 Hot Start High-Fidelity Polymerase (NEB). 5 μl of the resulting amplicon were diluted 1:4 in 1x buffer 2 (NEB), followed by denaturation and re-annealing in a nexus GSX1 Mastercycler (Eppendorf) by running the following protocol: Denaturation: 95 °C for 5 min; re-annealing: cooling down to 85 °C at a ramp rate of 0.2 °C/s followed by cooling down to 25 °C at a ramp rate of 0.1 °C/s. Next, 0.5 μl T7 endonuclease I (NEB) was added and the samples were incubated at 37 °C for 15 minutes. Samples were then analysed on 2 % Tris-borate-EDTA agarose gels. Gel documentation was performed using a Gel iX20 system equipped with a 2.8 megapixel/14 bit scientific grade CCD camera (INTAS). Intensities of DNA fragments were quantified using the ImageJ gel analysis tool (http://imagej.nih.gov/ij/). Finally, indel percentages were calculated using the following formula: indel (%) = 100x(1-(1-fraction cleaved)1/2), whereas the fraction cleaved = ∑(Cleavage product bands)/∑(Cleavage product bands + PCR input band). Full-length gel images are shown in Supplementary Figure S1.

### TIDE sequencing

Cells were lysed and genomic target loci were PCR amplified as described for the T7 assay. PCR amplicons were then resolved by gel electrophoresis followed by DNA isolation with a QIAquick Gel Extraction Kit (Qiagen). The amplicons were Sanger sequenced (Eurofins Genomics) and sequencing chromatograms were analysed using the TIDE web tool (https://tide.deskgen.com/) (52).

### Computational models of AcrIIC3-LOV2 hybrids

We used the Rosetta remodel application (53) to generate the AcrIIC3-LOV2 domain insertions based on the structures of AcrIIC3 (PDB 6J9N) and the LOV2 domain (PDB 2V0W). The N-terminus of the LOV2 structure contained three residues that were not part of our final design and thus omitted. Terminal regions of the LOV2 domain were rebuilt, including the added glycine-linkers. For rebuilding, fragment insertion with cyclic coordinate descent (54) and kinematic closure (55,56) with default parameters were used. For each of the variants, 1000 decoys were generated, of which 236 passed the chain-break filter for the AcrIIC3-LOV2 hybrid CN-C3G and 206 for the direct fusion CN-C3 (see below). These were subsequently clustered with a root mean square deviation threshold of 5 Å into 17 clusters for CN-C3G and 8 clusters for the direct fusion CN-C3.

### Statistical Analysis

Bars indicate means or single values, individual data points represent individual biological replicates, i.e. independent experiments performed on different days. For the luciferase experiments, each individual data point further represents the mean of three technical replicates, i.e. three separate wells of a 96-well plate transfected and treated in parallel. Error bars indicate the standard deviation (SD). Data analysis was performed with R (3.6.0).

## RESULTS

To create a photo-sensitive AcrIIC3 variant, we aimed at harnessing the LOV2 domain from *Avena sativa* (*As*) photoropin-1, which is a well-characterized conformational switch (Figure 1A). The *As*LOV2 domain carries two terminal helices denoted Jα and A’α, which, in the dark state, are docked against the LOV2 protein core so that the domain’s termini are in close proximity (distance ~10 Å) (57). Excitation with blue light triggers the unfolding and undocking of the Jα and A’α helices, resulting in a massive gain of flexibility at the LOV2 termini (58,59). It has previously been shown by Klaus Hahn and colleagues that inserting the *As*LOV2 domain into surface-exposed loops of enzymes can be used to disrupt their function in a light-dependent manner (60). We recently adapted this concept to engineer a light responsive variant of the *Spy*Cas9 inhibitor AcrIIA4 (29). By inserting the *As*LOV2 domain into the most C-terminal loop of AcrIIA4 (around residue E66/Y67) we created CASANOVA (for **C**RISPR-Cas9 **a**ctivity **s**witching via **a n**ovel **o**ptogenetic **v**ariant of **A**crIIA4), an AcrIIA4-LOV2 hybrid that blocks *Spy*Cas9 activity in the dark, but releases its function upon illumination. When developing CASANOVA, we relied on the available structural information, which guided our selection of LOV2 insertion sites on AcrIIA4. The AcrIIC3 structure, however, was not known at the start of this project and has only been reported very recently (48,61,62). Thus, we started this work by performing a secondary structure prediction using QUARK (63) with the goal to roughly identify regions corresponding to alpha-helices, beta-sheets and unstructured loops, as the latter are more permissive to domain insertions.

Based on this prediction, we chose seven target regions (R1-R7) in AcrIIC3 and inserted the LOV2 domain at several sites into each of these regions, most of which later turned out to correspond to actual loops (Figure 1B). The resulting AcrIIC3-LOV2 hybrids were then screened for their ability to block *Nme*Cas9 activity in the dark using a previously developed luciferase cleavage assay in HEK293T cells (49). In this assay, a catalytically active *Nme*Cas9 is targeted via a corresponding sgRNA to a firefly luciferase reporter gene (see Methods for details), thereby mediating strong reporter knockdown (Figure 1C, no Acr control). Co-supplying wild-type (wt) AcrIIC3 prevents reporter cleavage and thus results in a rescue of luciferase activity (Figure 1C, AcrIIC3). Remarkably, the AcrIIC3-LOV2 hybrids based on region 4 efficiently inhibited *Nme*Cas9 in the dark, as indicated by a potent rescue in luciferase activity comparable to that mediated by wt AcrIIC3 (Figure 1C). In contrast, the hybrids based on all other regions either showed considerably weakened inhibitory function as compared to wt AcrIIC3 (R7) or no *Nme*Cas9 inhibition at all (R1-3, R5-6). This suggests that LOV2 domain insertion at these sites interferes with Cas9 binding or impairs Acr folding.

To map the fusion site within R4 that would result in maximum inhibition in the dark and minimal inhibition upon illumination, we inserted the LOV2 domain behind each individual residue in this region, i.e. AcrIIC3 residues L58 to P64 (Supplementary Figure S2). Short deletions of up to three amino acids were optionally introduced at the target region, as such deletions turned out to be beneficial when engineering CASANOVA (29). Using the variants bearing the LOV2 domain behind AcrIIC3 residues F59 and N60 as scaffold (Supplementary Figure S2), we also tested the effects of inserting flexible GS-linkers at the Acr-LOV boundaries (Supplementary Figure S3). Finally, we optionally introduced the H519R/R521H double mutation into LOV2 (Supplementary Figure S3), as this was previously shown to improve the performance of a light-activated nuclear shuttle (LANS) based on LOV2 (64). We screened the resulting, comprehensive collection of AcrIIC3-LOV2 hybrids using the aforementioned luciferase assay and/or in genome editing experiments read out by T7 assay. To assess photosensitivity of the hybrid inhibitors, we either exposed samples to blue light for 48 hours (luciferase assays) or 72 hours (T7 assays) or kept them in the dark prior to measurements. Most AcrIIC3-LOV2 hybrids showed strong light-dependent Cas9 inhibition, albeit the level of Cas9 activity in the light and dark condition varied considerably between the different variants (Figure 1D-F, Supplementary Figures S2 and S3). The hybrids carrying the wt LOV2 domain behind AcrIIC3 residue F59 either with or without symmetric, single-glycine linkers at the LOV2-AcrIIC3 boundaries (variants 1 and 4 in Figure 1D, respectively) showed particularly robust light-switching behaviour in both assays, while blocking Cas9 activity in the dark comparably to wt AcrIIC3. We named these variants CN-C3 (for CASASNOVA-C3) and CN-C3G (for CASANOVA-C3 symmetric glycine linker variant), respectively.

Next, to investigate light-control when targeting *Nme*Cas9 to different, endogenous loci as well as upon different modes of delivery, HEK293T cells were either co-transfected with plasmids or co-transduced with AAV serotype-2 based vectors encoding (i) *Nme*Cas9 and a sgRNA and (ii) CN-C3(G). Indel formation at the target locus in the presence and absence of light was then measured by T7 assay and cross-validated by TIDE sequencing (Figure 2A).

**Figure 2.**
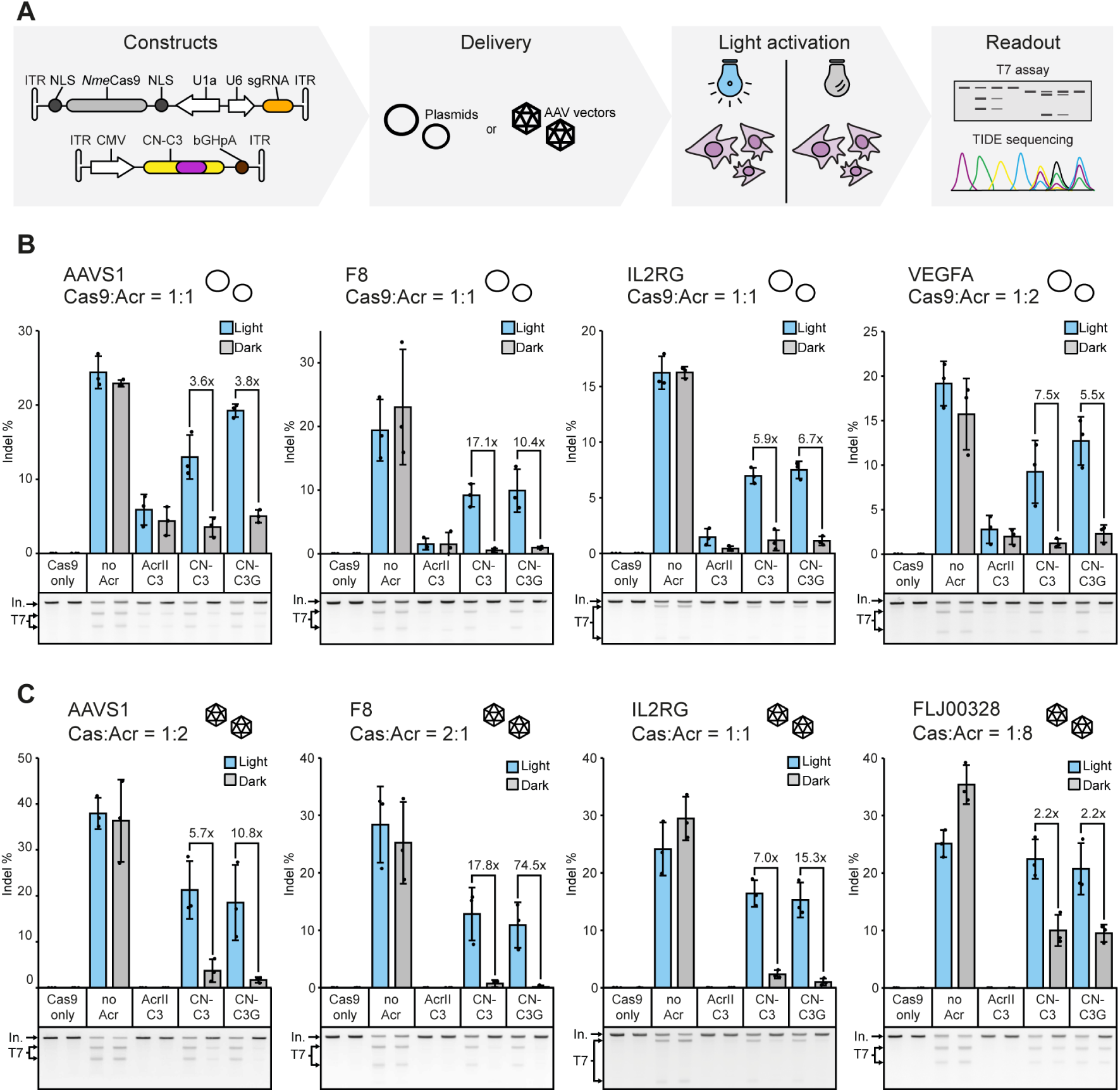
Light-dependent genome editing. (**A**) Experimental workflow. (**B, C**) HEK293T cells were co-transfected with plasmids (**B**) or co-transduced with AAV vectors (**C**) encoding (i) *Nme*Cas9 and a sgRNA targeting the indicated locus and (ii) the indicated Acr variant. Cells were then irradiated with pulsed blue light or kept in the dark for 72 hours, followed by T7 assay. Cas9:Acr vector mass ratios (**B**) and AAV lysate volume ratios (**C**) used during transfection or transduction, respectively, are indicated. Bars represent mean values, error bars the standard deviation and dots individual data points from n = 3 independent experiments. Representative gel images are shown below the bar charts. In, Input; T7, T7 cleavage fragments.

Remarkably, we observed potent, light-dependent editing at all loci tested (Figure 2B,C and Supplementary Figure S4). In line with previous observations in context of CASANOVA (29), the background editing in the dark was lower and the dynamic range of light-regulation slightly higher when applying AAV transduction instead of transient transfection for delivery (Figure 2C, Supplementary Figure S4), probably due to the more homogenous expression of the different components from AAVs. Interestingly, inhibition was to some degree locus-dependent, a property which was not specific to CN-C3(G), but also observed for wt AcrIIC3. When targeting the F8 locus, for instance, wt AcrIIC3 completely abolished indel formation, while for other loci, e.g. FLJ00328 considerable editing was observed also in presence of AcrIIC3. Similarly, our light-switchable CN-C3(G) system was extremely tight on some loci, while for other loci some editing also occurred in the absence of light. The performance of our *Nme*Cas9 light-switch was further dependent on the used CN-C3(G) dose. At increasing CN-C3(G):*Nme*Cas9 vector ratios, background editing in the dark was efficiently reduced, albeit at the cost of some reduction of Cas9 activity upon irradiation (Supplementary Figures S5 and S6). Together, these data demonstrate that our CN-C3 system is tuneable and can be used to efficiently control genome editing at various loci.

Having demonstrated optogenetic control of *Nme*Cas9 with CN-C3(G), we finally aimed at investigating possible mechanisms of LOV2-mediated light-switching. This was particularly relevant, as we did not use specific design criteria when engineering CN-C3(G) apart from trying to confine LOV2 insertion to loops (see above). First, we performed a detailed analysis of residue contacts within AcrIIC3 to see whether the loop into which we had inserted the LOV2 domain (region 4 in Figure 1B) connects interacting secondary structures. Unlike our previously reported LOV2 insertion site underlying CASANOVA (29) (Supplementary Figure S7), as well as those of previously published LOV2-kinase hybrids (60,65), the target loop within AcrIIC3 region 4 does not connect interacting secondary structures. Instead, it connects a helix and beta-sheet that stand in an angle of ~40 ° to one another (Figure 3A, B). Surprisingly, the insertion site appears to be located right at the boundary of the Cas9 binding surface (its distance to HNH domain is only ~7 Å) and, based on the reported structures of the AcrIIC3-*Nme*Cas9 HNH complex, would not even be considered entirely surface-exposed. In fact, the insertion site is directly flanked by residues, which mediate important contacts with the HNH domain (Figure 3B; AcrIIC3 L58 and N60 contacting Y540 and E539 on *Nme*Cas9, respectively). Of note, AcrIIC3 L58 is highly conserved and absolutely critical for AcrIIC3 function (61). Finally, to investigate possible configurations of the LOV2 domain in context of CN-C3(G), we performed structural modeling. In absence of an HNH binding partner, we found multiple possible conformations of the LOV2 domain relative to AcrIIC3 due to considerable flexibility at the Acr-LOV2 junction sites (Figure 3C and Supplementary Figure S8A). Out of the three most populated LOV2 conformational states, however, only one (cluster 3) did not show considerable steric clashes with the HNH domain when the CN-C3(G):HNH complex is assembled (Figure 3D, Supplementary Figure S8B, Supplementary Figure S9A). Structural alignment of the model to the recently published full-length *Nme*Cas9 bound to AcrIIC3 in 2:2 ratio (62) confirmed this result, suggesting that the position of the LOV2 domain relative to the Acr is restricted upon *Nme*Cas9 binding (Supplementary Figure S9B). Together, these observations suggest that the potent light-switching behaviour of CN-C3(G) might, at least in part, result from locally induced disorder directly at the Cas9 binding surface of AcrIIC3.

**Figure 3.**
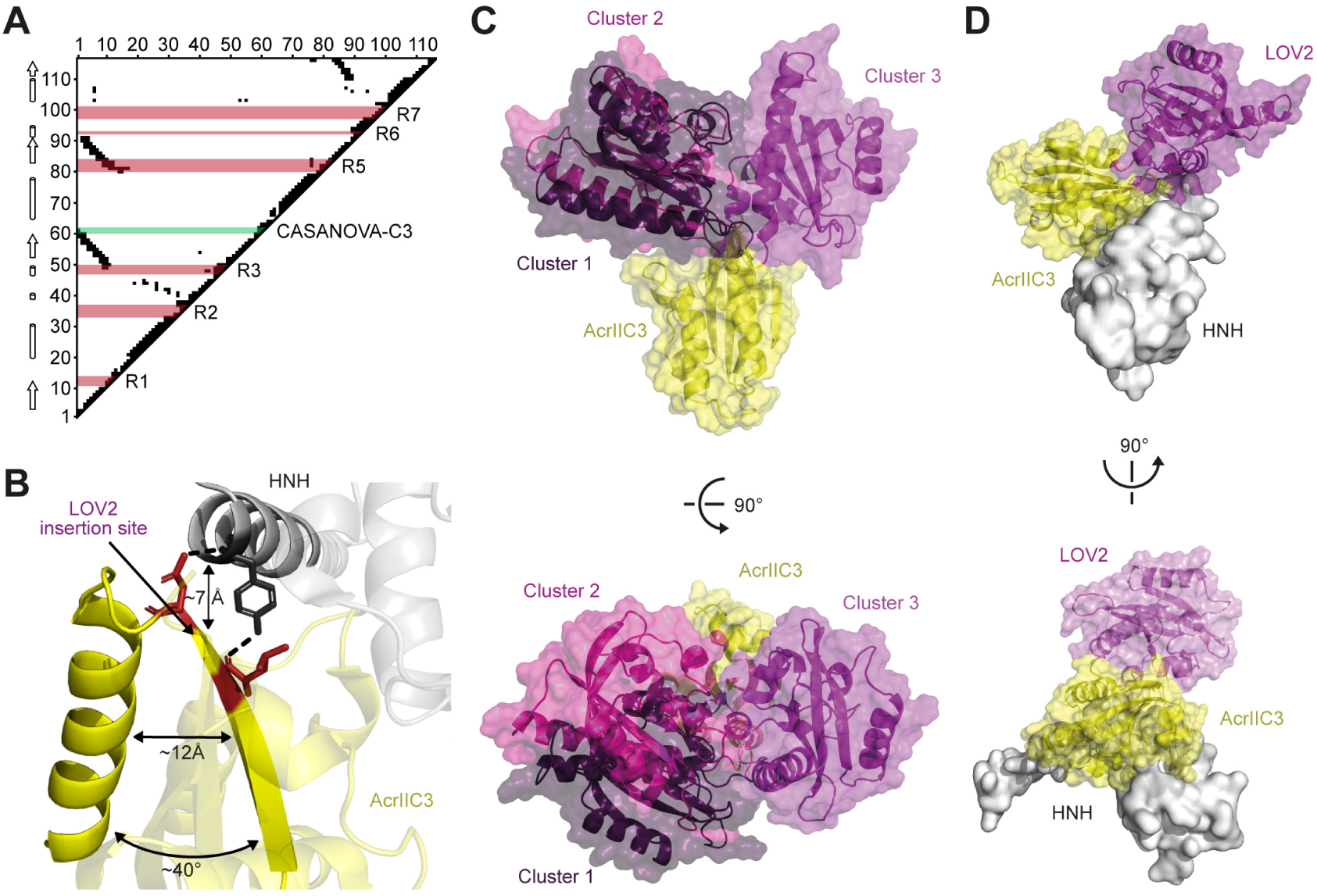
The LOV2 domain in CN-C3 is located in close proximity to the *Nme*Cas9 binding surface (**A**) Analysis of AcrIIC3 residue contacts. Spatially proximate AcrIIC3 residue pairs (distance <7 Å) are indicated by black squares. Secondary structure elements are shown on the left. Regions into which the LOV2 domain was inserted into AcrIIC3 (see Fig. 1B) are indicated in red. The LOV2 insertion site underlying CN-C3(G) is marked in green. Numbers correspond to AcrIIC3 residues. (**B**) Close-up view on the identified LOV2 insertion site in context of the AcrIIC3:HNH domain complex. The approximate distance between the insertion site on AcrIIC3 and the *Nme*Cas9 HNH domain is indicated. The angle as well as the distance between the secondary structure elements adjacent to the insertion site are shown. Residues in red mediate direct contact to the HNH domain. (**C**) Computational model of CN-C3 generated by domain assembly simulation. The three most populated conformational clusters of the LOV2 are shown in purple in descending order. (**D**) Cluster 3 does not sterically clash with the HNH-domain. PDB 6J9N, 2V0W.

## DISCUSSION

Systems to confine the activity of Cas9 in time and space are highly desired, as they improve the precision at which CRISPR-mediated genome perturbations can be made (25,26,30). We had previously engineered CASANOVA, an optogenetic *Spy*Cas9 inhibitor based on LOV2 insertion into AcrIIA4 (29). Here, we extended the CASANOVA approach to AcrIIC3, which is structurally unrelated to AcrIIA4, thereby demonstrating that LOV2 insertion into Acrs is a generalizable strategy to engineer light-switchable CRISPR inhibitors. CN-C3(G) enabled light-dependent *Nme*Cas9 genome editing at various target loci in mammalian cells and is compatible with delivery via transient transfection or AAVs. The latter are prime vector candidates for human gene therapy applications (66,67).

A particular advantage of light-switchable Acrs over photoactivatable Cas9 variants is their versatility: They are compatible with both, catalytically active Cas9 as well as dCas9-effector fusions, provided the underlying Acr impairs dCas9 DNA binding (as is the case for both, AcrIIA4 and AcrIIC3). Moreover, users can work with their established CRISPR constructs and systems such as Cas9 stable cell lines. We speculate that the future engineering of optogenetic Acrs based on broad-spectrum inhibitors such as AcrIIC1 (47) will further enhance their application range by enabling simultaneous regulation of multiple Cas9 orthologues.

In retrospect, in the absence of structural information of the Acr, we solely relied on sequence-based secondary structure predictions to guide our engineering efforts. Likely, the knowledge of the structure of the AcrIIC3:HNH complex would have lead us to exclude the best LOV2 domain insertion site due to its seemingly insufficient solvent-exposure and close proximity to the Cas9 binding surface. It is important to note that only few studies have been performed in the past, in which LOV2 domain insertion sites were mapped within target proteins in an unbiased fashion (68). Moreover, most past studies using LOV2 insertion for optogenetic regulation focused on enzymes, the engineering of which might follow different design criteria as compared to controlling protein-protein interactions as we do in this work. Thus, it will be interesting to explore whether LOV2 insertion in proximity to protein binding interfaces might be a generalizable design strategy for the engineering of light-dependent protein-protein interactions. Complementary, the unbiased mapping of LOV2 insertion sites within proteins of different origin and function might be a very interesting strategy to obtain new and powerful, yet unconventional LOV2-hybrid designs for optogenetic applications. Together, our work yielded the first tool for optogenetic control of *Nme*Cas9-mediated genome editing and suggests a novel approach to engineer light-dependent protein-protein interactions.

## AVAILABILITY

The CN-C3 and CN-C3G vectors will be made available via Addgene. Annotated plasmid sequences (SnapGene DNA files) are provided as Supplementary Data. Structural models of CN-C3(G) are also available as Supplementary Data. All other data is available from the corresponding authors on reasonable request. Code and data for the models of CN-C3(G) will be made available on GitHub (https://github.com/juzb/CASANOVA-C3).

## SUPPLEMENTARY DATA

Supplementary Figures 1-9 and Supplementary Tables 1-3 are provided as Supplementary Data.

## ACKNOWLEDGEMENT

We thank the Synthetic Biology group (Heidelberg University hospital) for their support and critical feedback on the manuscript. We are thankful to Erik J. Sontheimer (University of Massachusetts Medical School, Worcester) for sharing a plasmid.

## Author contributions

D.N. conceived the study. M.D.H., J.M., C.S. and D.N. designed experiments. M.D.H. and J.M. conducted experiments. M.D.H., J.M. and D.N. analysed and interpreted data. J.U.z.B and Z.H. performed computational modeling of the AcrIIC3-LOV2 hybrid structures. R.E. and D.N. jointly directed the work with support from B.E.C.. J.M. and D.N. wrote the manuscript with support from all authors.

