## Supplementary Information for "Optogenetic control of *Neisseria meningitidis* Cas9 genome editing using an engineered, light-switchable anti-CRISPR protein"

Figure 1F



Figure 2B

AAVS1 F8 IL2RG VEGFA

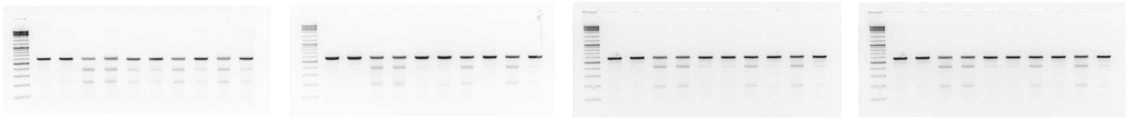

Figure 2C

AAVS1 F8 IL2RG FLJ00328

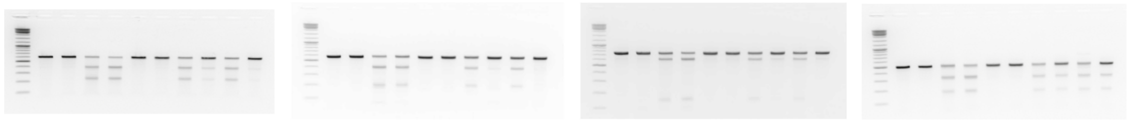

Supplementary Figure S2

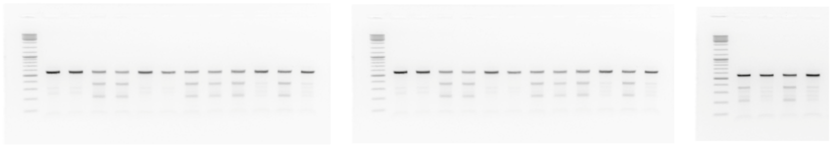

Supplementary Figure S5

Cas9:Acr 2:1 1:1 1:2

AAVS1

F8

IL2RG

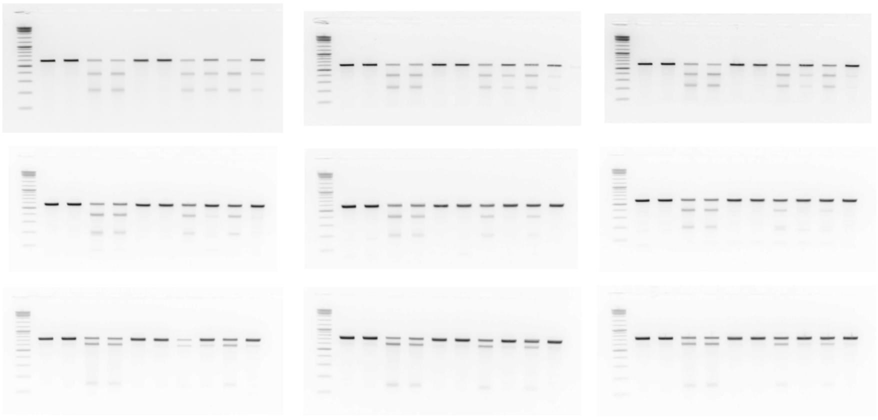

**Supplementary Figure 1.** Full length gel images. The Gene Ruler DNA Ladder Mix (Thermo Fisher Scientific) was used in all cases (left lane).

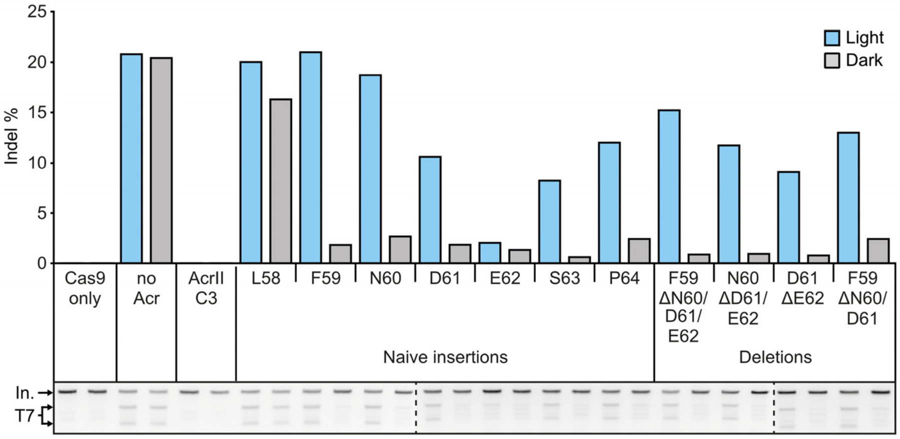


**Supplementary Figure 2.** Systematic screen for LOV2 insertion sites within region 4. HEK293T cells were transfected with plasmids encoding (i) *Nme*Cas9 and an IL2RG locus targeting sgRNA and (ii) the indicated AcrIIC3 variant in a vector mass ratio of 1:2. Samples were either exposed to blue light or kept in the dark for 72 hours followed by T7 assay. AcrIIC3 residues behind which the LOV2 domain was inserted as well as residues deleted from AcrIIC3 are indicated. Bars represent indel frequencies. Gel images are shown below the bars. The dotted lines separate different gels. Data correspond to a single experiment. In, Input; T7, T7 cleavage fragments.

**A**

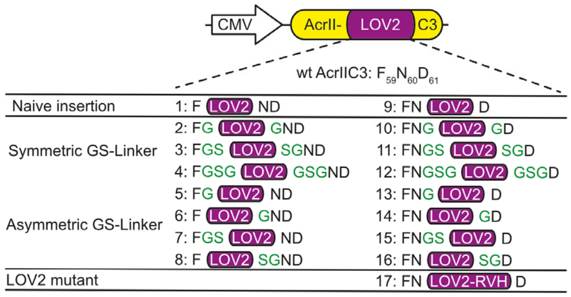


**B**

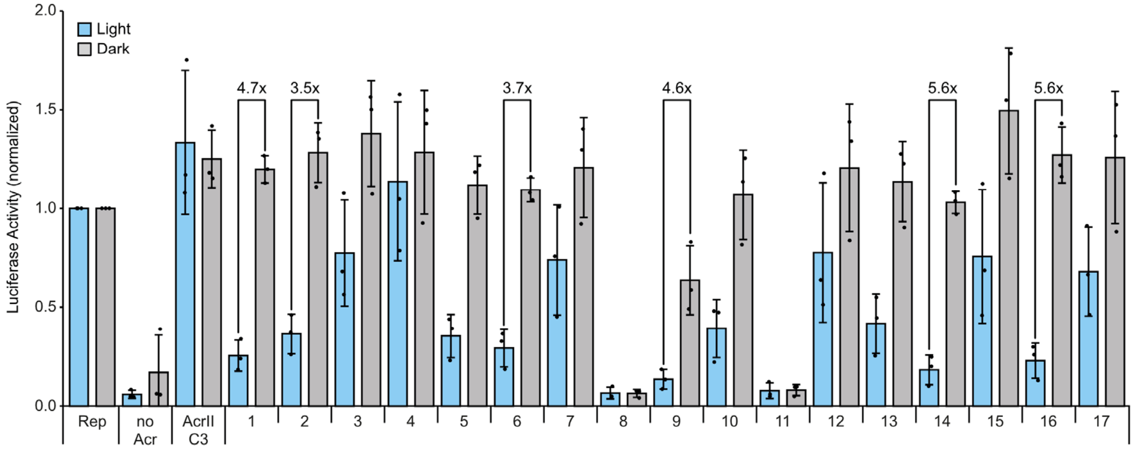


**Supplementary Figure 3.** Optimization of AcrIIC3-LOV2 hybrids by linker insertion and LOV2 domain mutation. (**A**) A library of 17 constructs was designed using the variants with the LOV2 domain inserted behind F59 or N60 as scaffold. Short symmetric and asymmetric glycine-serine linkers were included at the Acr-LOV boundaries (indicated in green). Variant 17 carries a LOV2 H519R/R521H double mutant. (**B**) Luciferase assay-based screening of the variants in **A**. HEK293T cells were transfected with constructs encoding (i) a luciferase reporter, (ii) *Nme*Cas9 and a sgRNA targeting the reporter gene and (iii) the indicated AcrIIC3-LOV2 hybrid. Samples were irradiated with blue light or kept in the dark for 48 hours. Bars represent means, error bars the standard deviation and dots individual data points from n = 3 independent experiments. Rep, reporter only control.

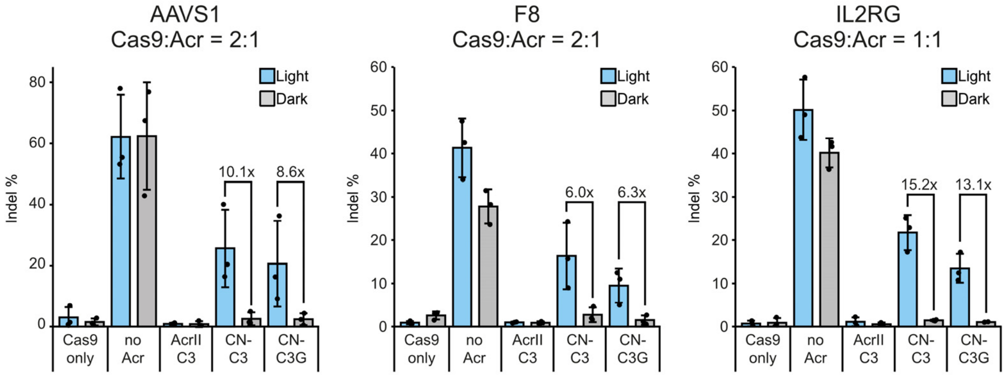


**Supplementary Figure 4.** Light-dependent genome editing using AcrIIC3-LOV2 hybrids assessed by TIDE sequencing. HEK293T cells were co-transduced with AAVs encoding (i) *Nme*Cas9 and the indicated sgRNA and (ii) CN-C3(G) or wt AcrIIC3 in the indicated AAV lysate volume ratios. Indel frequencies were assessed by TIDE sequencing. Bars represent means, error bars the standard deviation and dots individual data points from n = 3 independent experiments.

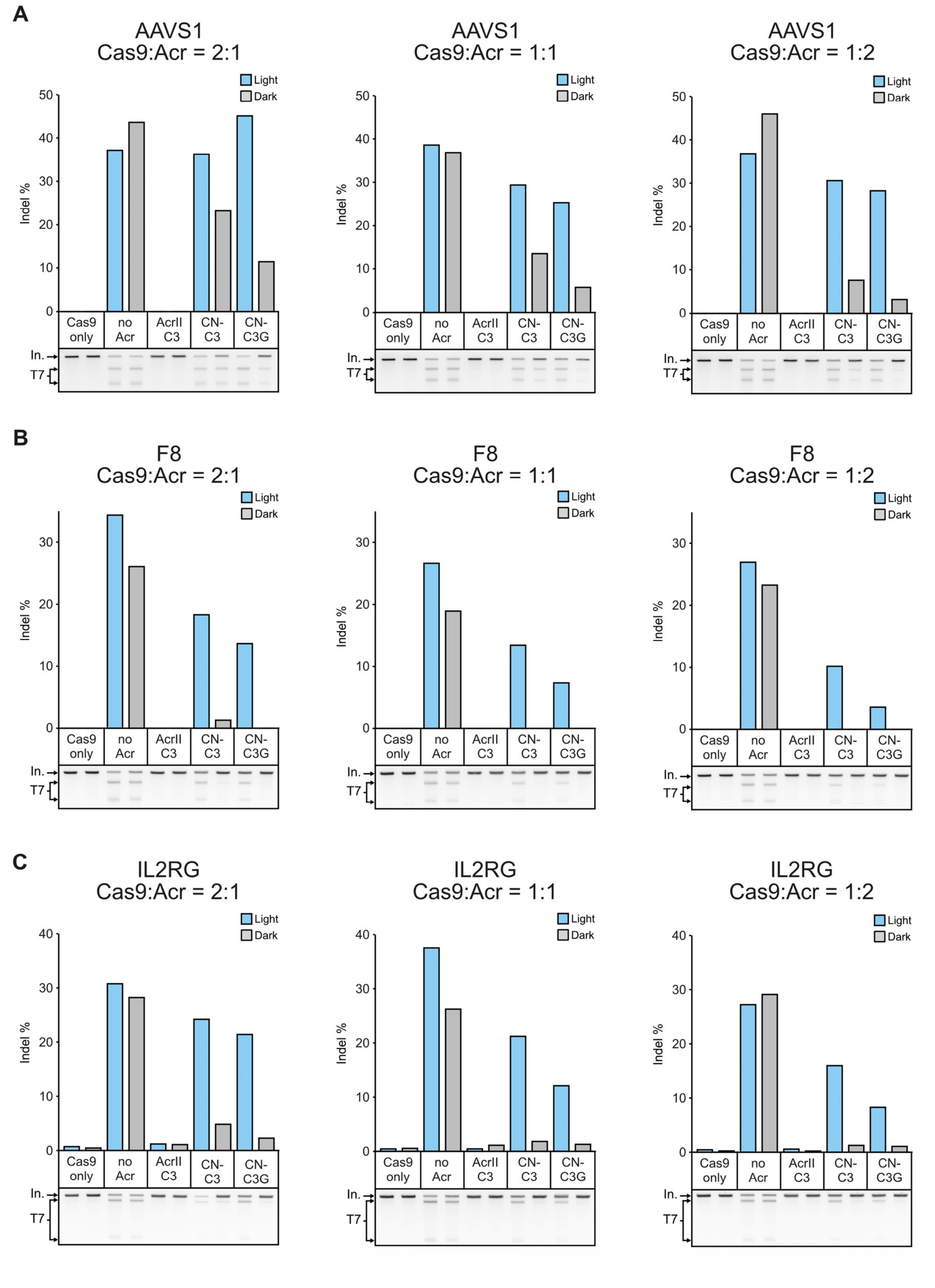


**Supplementary Figure 5.** *Nme*Cas9 photo-regulation depends on the used CN-C3(G) AAV vector dose. HEK293T cells were co-transduced with AAV vectors encoding (i) *Nme*Cas9 and a sgRNA targeting the AAVS1 (**A**), F8 (**B**) or IL2RG (**C**) locus and (ii) the respective Acr in the indicated AAV lysate volume ratios. Indel frequencies were analyzed by T7 assay. Bars represent indel frequencies as quantified from the gel images shown below the bars. Data correspond to a single experiment. In, Input; T7, T7 cleavage fragments.

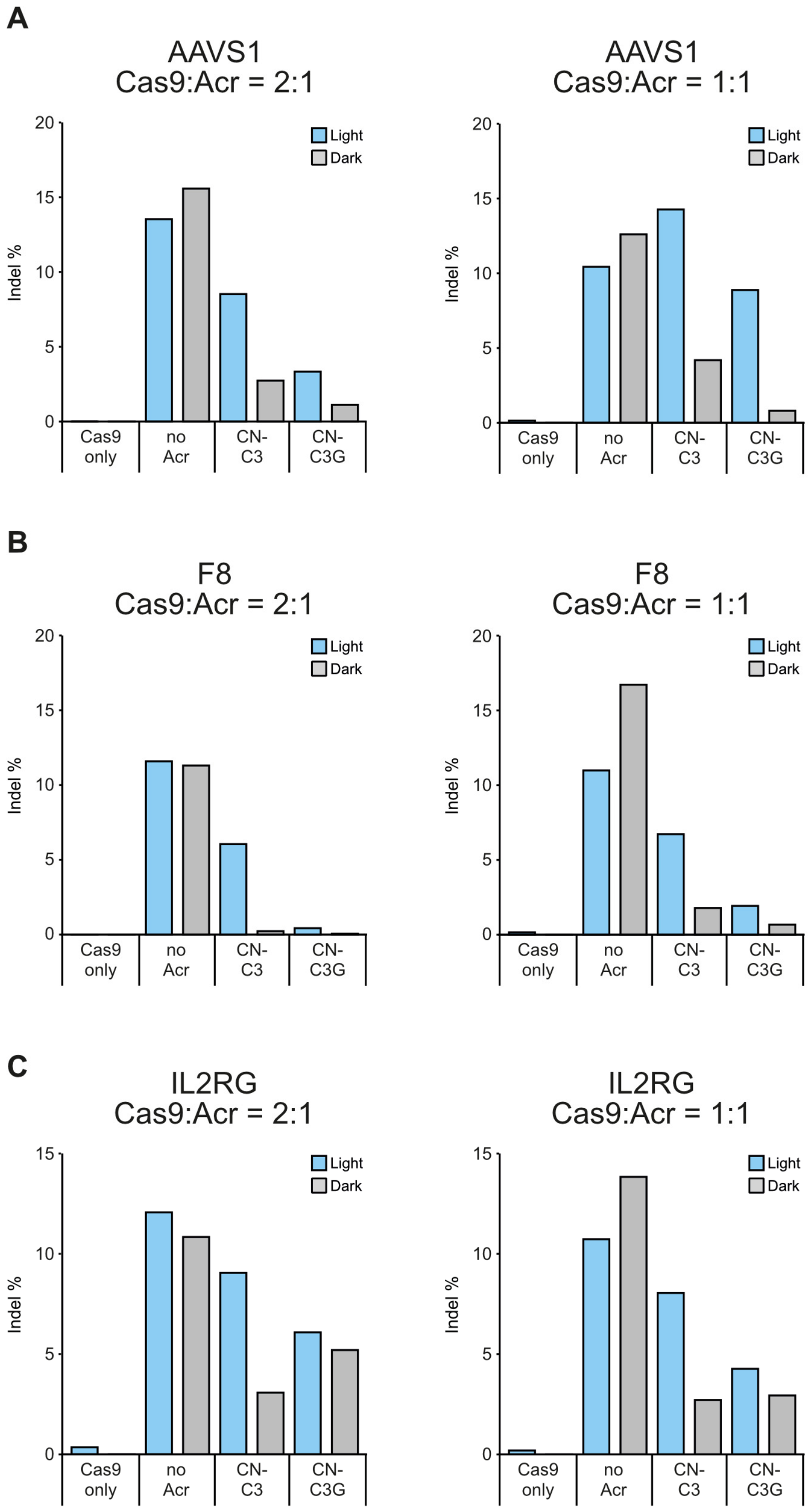


**Supplementary Figure 6.** *Nme*Cas9 photo-regulation depends on the CN-C3(G) vector dose used during transfection. HEK293T cells were co-transfected with constructs encoding (i) *Nme*Cas9 and a sgRNA targeting the AAVS1 (**A**), F8 (**B**) or IL2RG (**C**) locus and (ii) CN-C3(G) or wt AcrIIC3 using the indicated vector mass ratios. Indel frequencies were assessed by T7 assay. Bars represent indel frequencies corresponding to a single experiment.

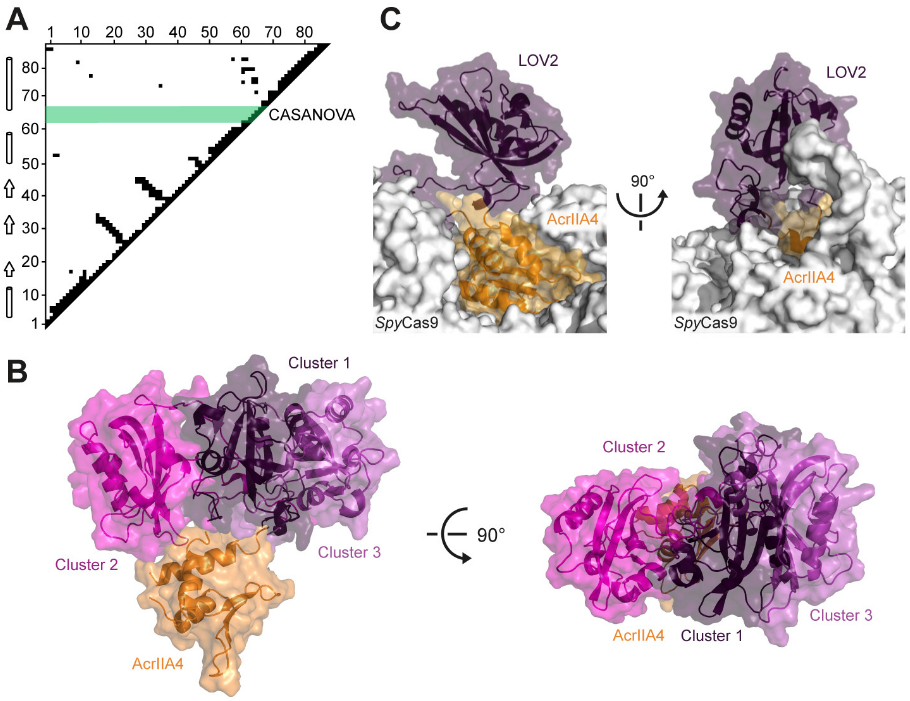


**Supplementary Figure 7.** Structural analysis of CASANOVA, an optogenetic *Spy*Cas9 inhibitor. The data underlying this figure has been previously reported by us (1) and is shown here to facilitate comparison with CN-C3(G) (Figure 3 and Supplementary Figure S8). (**A**) Analysis of AcrIIA4 residue contacts. Spatially proximate AcrIIA4 residue pairs (distance <7 Å) are indicated by black squares. Secondary structure elements are shown on the left. The region, in which the LOV2 domain was inserted into AcrIIC3 is marked in green. (**B**) Computational model of CASANOVA generated by domain assembly simulation. The three most populated LOV2 conformational clusters are shown in purple in descending order. (**C**) Structural model of CASANOVA bound to *Spy*Cas9. The shown LOV2 configuration corresponds to the cluster 1 in C. (**B, C**) PDB 5VW1 and 2V0W.

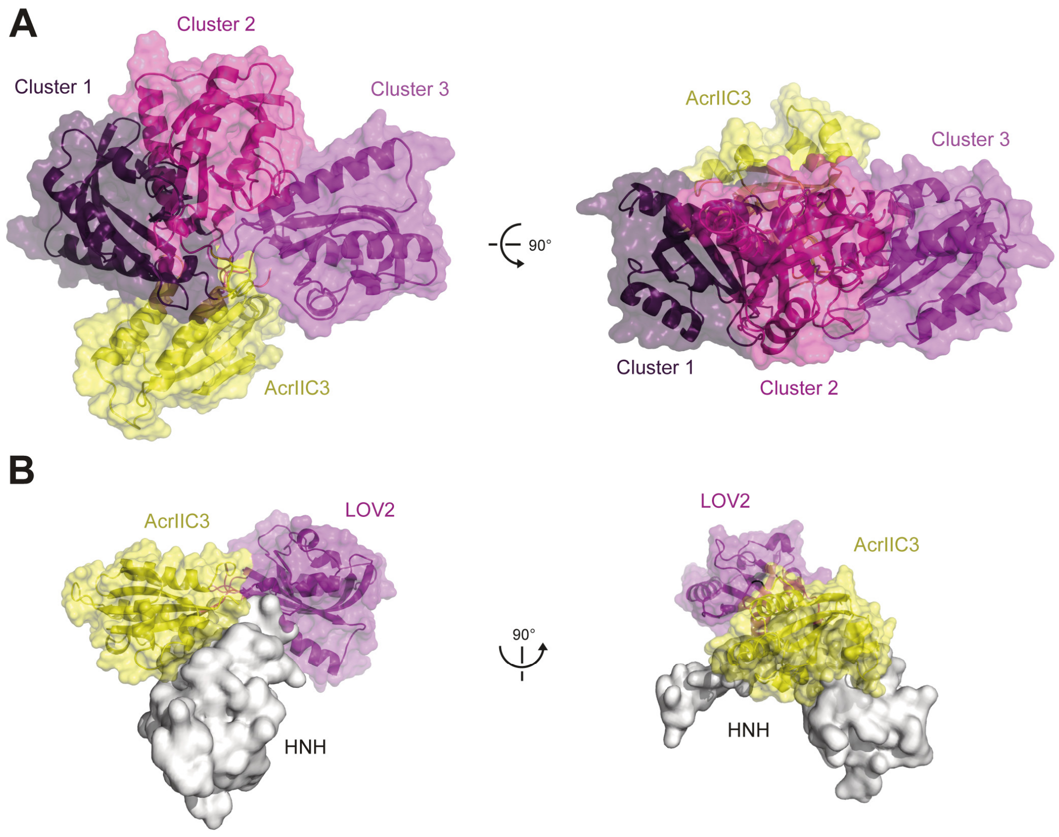


**Supplementary Figure 8.** Structural models of CN-C3G. (**A**) The three most populated LOV2 conformational clusters as generated by domain assembly simulations are shown. (**B**) In complex with the HNH domain of *Nme*Cas9, only cluster 3 does not show steric clashes. PDB 6J9N, 2V0W.

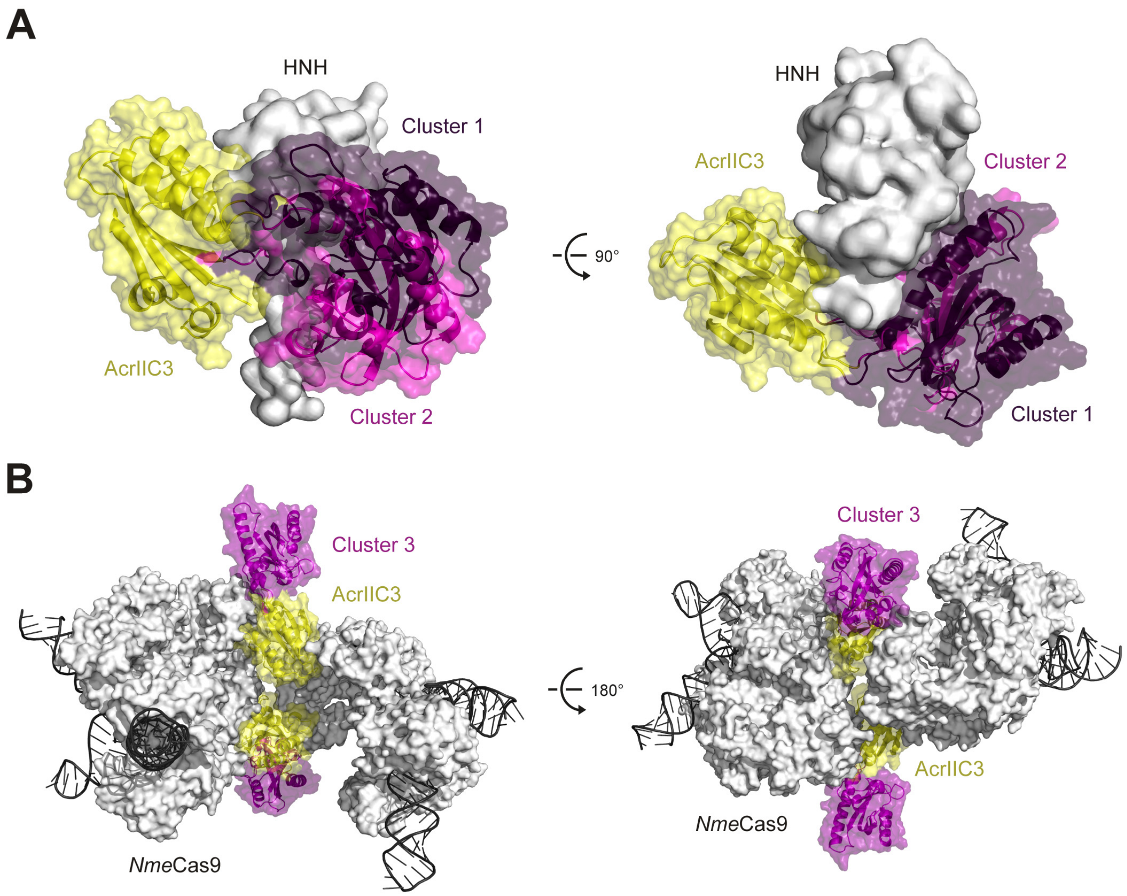


**Supplementary Figure 9.** (**A**) Two of the three most populated LOV2 conformational clusters sterically clash with the HNH domain. The two most populated cluster of CN-C3 in the HNH-bound form are shown. PDB 6J9N and 2V0W. (**B**) Alignment of the CN-C3 model with LOV2 cluster 3 to the structure of the *Nme*Cas9-AcrIIC3 dimeric complex (2) (PDB 6JE9).

**Supplementary Table 1.** List of constructs. CMV, cytomegalo virus; AAV, adeno-associated virus; SV40, Simian Virus 40; TK, thymidine kinase; NLS, nuclear localization signal; HA, human influenca hemagglutinin.

| # | Name | Description | Source |
|---|---|---|---|
| 1 | Dual luciferase reporter | SV40 promoter firefly luciferase; TK promoter *Renilla* luciferase; sgRNA targeting firefly gene | (3) |
| 2 | h*Nme*Cas9 + sgRNA scaffold (pEJS654 All-in-One AAV-sgRNA-h*Nme*Cas9; Addgene plasmid: #112139) | U1a promoter NLS h*Nme*Cas9 NLS 3xHA; U6 promoter sgRNA scaffold | (4) |
| 3 | h*Nme*Cas9 + VEGFA sgRNA (AAV) | U1a promoter NLS h*Nme*Cas9 NLS 3xHA; U6 promoter VEGFA sgRNA | (5) |
| 4 | h*Nme*Cas9 + IL2RG sgRNA (AAV) | U1a promoter NLS h*Nme*Cas9 NLS 3xHA; U6 promoter IL2RG sgRNA | (3) |
| 5 | h*Nme*Cas9 + FLJ00328 sgRNA (AAV) | U1a promoter NLS h*Nme*Cas9 NLS 3xHA; U6 promoter FLJ00328 sgRNA | (3) |
| 6 | h*Nme*Cas9 + AAVS1 sgRNA (AAV) | U1a promoter NLS h*Nme*Cas9 NLS 3xHA; U6 promoter AAVS1 sgRNA | (3) |
| 7 | h*Nme*Cas9 + F8 sgRNA (AAV) | U1a promoter NLS h*Nme*Cas9 NLS 3xHA; U6 promoter F8 sgRNA | (3) |
| 8 | Wild-type AcrIIC3 | CMV promoter AcrIIC3 | (5) |
| 9 | AcrIIC3 S11-LOV2-F12 | CMV promoter AcrIIC3 S11-LOV2-F12 | This work |
| 10 | AcrIIC3 F12-LOV2-N13 | CMV promoter AcrIIC3 F12-LOV2-N13 | This work |
| 11 | AcrIIC3 N13-LOV2-G14 | CMV promoter AcrIIC3 N13-LOV2-G14 | This work |
| 12 | AcrIIC3 R33-LOV2-V34 | CMV promoter AcrIIC3 R33-LOV2-V34 | This work |
| 13 | AcrIIC3 V34-LOV2-S35 | CMV promoter AcrIIC3 V34-LOV2-S35 | This work |
| 14 | AcrIIC3 S35-LOV2-I36 | CMV promoter AcrIIC3 S35-LOV2-I36 | This work |
| 15 | AcrIIC3 I36-LOV2-I37 | CMV promoter AcrIIC3 I36-LOV2-I37 | This work |
| 16 | AcrIIC3 N47-LOV2-A48 | CMV promoter AcrIIC3 N47-LOV2-A48 | This work |
| 17 | AcrIIC3 A48-LOV2-S49 | CMV promoter AcrIIC3 A48-LOV2-S49 | This work |
| 18 | AcrIIC3 S49-LOV2-L50 | CMV promoter AcrIIC3 S49-LOV2-L50 | This work |
| 19 | AcrIIC3 N60-LOV2-D61 | CMV promoter AcrIIC3 N60-LOV2-D61 | This work |
| 20 | AcrIIC3 D61-LOV2-E62 | CMV promoter AcrIIC3 D61-LOV2-E62 | This work |
| 21 | AcrIIC3 T81-LOV2-G82 | CMV promoter AcrIIC3 T81-LOV2-G82 | This work |
| 22 | AcrIIC3 I83-LOV2-S84 | CMV promoter AcrIIC3 I83-LOV2-S84 | This work |
| 23 | AcrIIC3 E92-LOV-S93 | CMV promoter AcrIIC3 E92-LOV-S93 | This work |
| 24 | AcrIIC3 R97-LOV2-L98 | CMV promoter AcrIIC3 R97-LOV2-L98 | This work |
| 25 | AcrIIC3 L98-LOV2-P99 | CMV promoter AcrIIC3 L98-LOV2-P99 | This work |
| 26 | AcrIIC3 P99-LOV2-V100 | CMV promoter AcrIIC3 P99-LOV2-V100 | This work |
| 27 | AcrIIC3 V100-LOV2-E101 | CMV promoter AcrIIC3 V100-LOV2-E101 | This work |
| 28 | AcrIIC3 L58-LOV2-F59 | CMV promoter AcrIIC3 L58-LOV2-F59 | This work |
| 29 | AcrIIC3 F59-LOV2-N60 (**CN-C3**) | CMV promoter AcrIIC3 F59-LOV2-N60 | This work |
| 30 | AcrIIC3 N60-LOV2-D61 | CMV promoter AcrIIC3 N60-LOV2-D61 | This work |
| 31 | AcrIIC3 D61-LOV2-E62 | CMV promoter AcrIIC3 D61-LOV2-E62 | This work |
| 32 | AcrIIC3 E62-LOV2-S63 | CMV promoter AcrIIC3 E62-LOV2-S63 | This work |
| 33 | AcrIIC3 S63-LOV2-P64 | CMV promoter AcrIIC3 S63-LOV2-P64 | This work |
| 34 | AcrIIC3 P64-LOV2-A65 | CMV promoter AcrIIC3 P64-LOV2-A65 | This work |

| | | | |
|---|---|---|---|
| 35 | AcrIIC3 F59-LOV2-S63 | CMV promoter AcrIIC3 F59-LOV2-S63 | This work |
| 36 | AcrIIC3 N60-LOV2-S63 | CMV promoter AcrIIC3 N60-LOV2-S63 | This work |
| 37 | AcrIIC3 D61-LOV2-S63 | CMV promoter AcrIIC3 D61-LOV2-S63 | This work |
| 38 | AcrIIC3 F59-LOV2-E62 | CMV promoter AcrIIC3 F59-LOV2-E62 | This work |
| 39 | AcrIIC3 F59-G-LOV2-G-N60 (**CN-C3G**) | CMV promoter AcrIIC3 F59-G-LOV2-G-N60 | This work |
| 40 | AcrIIC3 F59-GS-LOV2-SG-N60 | CMV promoter AcrIIC3 F59-GS-LOV2-SG-N60 | This work |
| 41 | AcrIIC3 F59-GSG-LOV2-GSG-N60 | CMV promoter AcrIIC3 F59-GSG-LOV2-GSG-N60 | This work |
| 42 | AcrIIC3 F59-G-LOV2-N60 | CMV promoter AcrIIC3 F59-G-LOV2-N60 | This work |
| 43 | AcrIIC3 F59-LOV2-G-N60 | CMV promoter AcrIIC3 F59-LOV2-G-N60 | This work |
| 44 | AcrIIC3 F59-GS-LOV2-N60 | CMV promoter AcrIIC3 F59-GS-LOV2-N60 | This work |
| 45 | AcrIIC3 F59-LOV2-SG-N60 | CMV promoter AcrIIC3 F59-LOV2-SG-N60 | This work |
| 46 | AcrIIC3 N60-G-LOV2-G-D61 | CMV promoter AcrIIC3 N60-G-LOV2-G-D61 | This work |
| 47 | AcrIIC3 N60-GS-LOV2-SG-D61 | CMV promoter AcrIIC3 N60-GS-LOV2-SG-D61 | This work |
| 48 | AcrIIC3 N60-GSG-LOV2-GSG-D61 | CMV promoter AcrIIC3 N60-GSG-LOV2-GSG-D61 | This work |
| 49 | AcrIIC3 N60-G-LOV2-D61 | CMV promoter AcrIIC3 N60-G-LOV2-D61 | This work |
| 50 | AcrIIC3 N60-LOV2-G-D61 | CMV promoter AcrIIC3 N60-LOV2-G-D61 | This work |
| 51 | AcrIIC3 N60-GS-LOV2-D61 | CMV promoter AcrIIC3 N60-GS-LOV2-D61 | This work |
| 52 | AcrIIC3 N60-LOV2-SG-D61 | CMV promoter AcrIIC3 N60-LOV2-SG-D61 | This work |
| 53 | AcrIIC3 D61-G-LOV2-S63 | CMV promoter AcrIIC3 D61-G-LOV2-S63 | This work |
| 54 | AcrIIC3 LOV2-RVH_N60D61 | CMV promoter AcrIIC3 LOV2-RVH_N60D61 | This work |
| 55 | AAV AcrIIC3 (Addgene: # 120301) | AAV compatible vector encoding AcrIIC3 | (5) |
| 56 | AAV AcrIIC3 F59-LOV2-N60 | AAV compatible vector encoding AcrIIC3 F59-LOV2-N60 | This work |
| 57 | AAV AcrIIC3 F59-G-LOV2-G-N60 | AAV compatible vector encoding AcrIIC3 F59-G-LOV2-G-N60 | This work |
| 58 | pBluescript | Empty vector | Invitrogen |

**Supplementary Table 2.** Genomic target sites used for genome editing experiments. Spacer sequences are underlined. PAM motifs are in bold.

| Locus | Sequence 5' –> 3' |
|---|---|
| IL2RG (6) | CTCTTTCTCCTCAAGGAACAATCAGTG**GATT** |
| FLJ00328 (6) | GGACAGGAGTCGCCAGAGGCCGGTGGTG**GATT** |
| AAVS1 (6) | ACCCCACAGTGGGGCCACTAGGGACAG**GATT** |
| F8 (6) | GGTTTCTAGTTGTGACAAGAACACTGGT**GATT** |
| VEGFA (6) | GCGGGGAGAAGGCCAGGGGTCACTCCAG**GATT** |

**Supplementary Table 3.** List of primers used for genomic PCRs for T7 assay and TIDE sequencing.

| Locus | Direction | Sequence 5' –> 3' |
|---|---|---|
| IL2RG | Forward | ATGACACTGGTGGGTGTTCAG |
| | Reverse | TCTTCACCTTGCAGGCTCTCT |
| FLJ00328 | Forward | AGAGGAGCCTTCTGACTGCTGCAGA |
| | Reverse | AGGTCCTGGCCTTGCCTTCGA |
| AAVS1 | Forward | TGCTTTCTTTGCCTGGACAC |
| | Reverse | CCTCTCTGGCTCCATCGTAA |
| F8 | Forward | GGGAGAGAACCTCTAACAGAACG |
| | Reverse | GCTCCAGGTGATGGATCATCAG |
| VEGFA | Forward | GTGTGCAGACGGCAGTCACTAG |
| | Reverse | CTCTGCGGACGCTCAGTGAAG |
